# Homologous recombination deficiency prediction from whole slide images using label refinement and foundation-model benchmarking in ovarian cancer

**DOI:** 10.64898/2026.06.25.734452

**Authors:** Nauman Ali Shah, Makhdoom Sarwar, Ehsan Ullah

**Affiliations:** Senior Analytics Specialist, Westpac New Zealand; Research Fellow, Department of Gynaecology and Obstetrics, Otago University, New Zealand; Adjunct Assistant Professor, Department of Pathology, The Ohio State University, Columbus, OH, USA; Director, Translyx Limited, Auckland, New Zealand; Service Manager, Surgical Division, Health New Zealand, Counties Manukau

**Keywords:** Computational pathology, digital pathology, ovarian cancer, high-grade serous ovarian carcinoma, homologous recombination deficiency, HRD, PARP inhibitor, whole-slide imaging, foundation models, TCGA-OV, multiple instance learning, label refinement

## Abstract

**Background:** Homologous recombination deficiency (HRD) is clinically imperative in high-grade serous ovarian carcinoma (HGSOC), particularly because of its association with platinum sensitivity and benefit from poly(ADP-ribose) polymerase inhibitor (PARPi) therapy. However, public datasets rarely contain a complete combination of diagnostic haematoxylin and eosin (H&E) whole-slide images (WSIs), validated clinical HRD assay results, genomic scar scores, BRCA1 promoter methylation data, and treatment-response outcomes. This creates a major barrier for computational pathology studies seeking to develop clinically interpretable models of HRD or PARPi response from routine histology.

**Objective:** We performed an exploratory, leakage-controlled computational pathology benchmarking study to evaluate whether H&E WSIs from TCGA-OV contain a measurable morphology-linked signal associated with research-grade molecular HRD labels, and whether label refinement and pathology foundation-model embeddings alter predictive performance.

**Methods:** We assembled a frozen-primary TCGA-OV WSI cohort comprising 717 tissue-section/biospecimen slides from 316 patients. Diagnostic FFPE DX slides were excluded from model selection because of complete patient overlap with the frozen-primary cohort. Two HRD labels were evaluated: an initial mutation-only molecular label based on BRCA/HR-gene mutation evidence, and a refined methylation-enhanced molecular label that additionally incorporated BRCA1 promoter methylation. Feature extraction was performed using ResNet50, UNI, CONCH, Virchow2, Phikon-v2, and UNI2-h encoders. Patient-level attention-based multiple instance learning (ABMIL) was used with patient-as-bag modelling. Evaluation used patient-level grouped 5-fold x 5-repeat stratified cross-validation, with 25 folds total, bootstrap confidence intervals, and patient-level leakage control.

**Results:** The initial mutation-only label classified 78 patients as positive and 238 as negative. The refined methylation-enhanced label recovered 33 additional positives, resulting in 111 positive and 205 negative patients. Patient-level ABMIL using UNI2-h features achieved the strongest performance for the refined label, with AUROC 0.634 (95% CI 0.571-0.698), AUPRC 0.468 (95% CI 0.390-0.562), balanced accuracy 0.597, sensitivity 0.532, specificity 0.663, F1 score 0.494, and Brier score 0.233. The calibrated threshold was 0.512, yielding TN=136, FP=69, FN=52, and TP=59. Comparative models showed lower discrimination, including UNI2-h with the initial label (AUROC 0.628), Phikon-v2 refined (0.582), Virchow2 refined (0.582), CONCH initial (0.587), ResNet50 refined (0.570), and clinical baselines (AUROC 0.54-0.57).

**Conclusions:** TCGA-OV H&E WSIs contain a modest but reproducible morphology-linked signal associated with research-grade molecular HRD status. However, the AUROC around 0.63, absence of clinical HRD assay labels, lack of genomic scar endpoints in the implemented workflow, and absence of PARPi/platinum response targets prevent clinical interpretation. This study should be interpreted as a proof-of-concept benchmarking framework and methodological foundation for future H&E-based predictive modelling in clinically curated PARPi response cohorts.

## INTRODUCTION

High-grade serous ovarian carcinoma (HGSOC) is the most common and lethal subtype of epithelial ovarian cancer and is characterised by extensive genomic instability, near-universal TP53 mutation, frequent copy-number alteration, and a clinically important subset of tumours with homologous recombination deficiency (HRD).^1–4^ HRD reflects impaired repair of DNA double-strand breaks through homologous recombination repair pathways and is associated with sensitivity to platinum chemotherapy and poly(ADP-ribose) polymerase inhibitor (PARPi) therapy.^5–12^ The clinical relevance of HRD has been reinforced by major PARPi maintenance trials in ovarian cancer, including SOLO1, PAOLA-1, PRIMA, NOVA, and ARIEL3.^13–15^

Despite this clinical importance, HRD is not a single molecular entity. It may arise through germline or somatic BRCA1/2 alterations, BRCA1 promoter methylation, or defects in other homologous recombination repair genes.^16–19^ It may also be inferred through genomic scar assays based on loss of heterozygosity (LOH), telomeric allelic imbalance (TAI), and large-scale state transitions (LST).^20–25^ These biomarkers capture different aspects of HRD biology and vary in availability across datasets and clinical settings. Public datasets such as The Cancer Genome Atlas ovarian carcinoma cohort (TCGA-OV) provide invaluable molecular and histopathological data, but they do not uniformly provide the full set of information needed to build a clinically validated HRD or PARPi-response model from H&E slides.^1,26^

Computational pathology has shown that routine H&E WSIs can encode information associated with diagnosis, prognosis, molecular alterations, tumour microenvironment, and treatment response.^27–35^ Multiple instance learning (MIL) and attention-based MIL approaches have become widely used for WSI-level prediction because they allow slide-level or patient-level labels to be learned from gigapixel images without requiring exhaustive pixel-level annotation.^36–39^ More recently, pathology foundation models such as UNI, CONCH, Virchow, and other large-scale histology encoders have enabled transfer learning from broad histopathology pretraining to downstream disease-specific tasks.^40–44^ These developments raise the possibility that routine H&E morphology may contain measurable signals associated with HRD biology in HGSOC.

However, H&E-based HRD prediction remains challenging. Recent studies, including DeepHRD and ovarian cancer-specific deep-learning analyses, suggest that HRD or BRCA-related states may be partly inferable from histology.^45–48^ At the same time, the field remains vulnerable to label noise, slide-level leakage, dataset bias, slide-type confounding, and overstatement of clinical claims when public molecular proxies are used as substitutes for validated clinical endpoints.^49–53^ A rigorous benchmarking study must therefore explicitly separate research-grade proxy modelling from clinical HRD or PARPi-response prediction.

In this study, we developed and evaluated a leakage-controlled computational pathology pipeline for HRD-molecular label prediction from H&E WSIs in TCGA-OV. We tested two central hypotheses: first, that H&E morphology may contain a modest but measurable signal associated with research-grade molecular HRD labels; and second, that label refinement incorporating BRCA1 promoter methylation may improve biological plausibility and alter apparent model performance. We compared conventional and pathology foundation-model feature extractors, including ResNet50, UNI, CONCH, Virchow2, Phikon-v2 and UNI2-h, using patient-level attention-based MIL and grouped cross-validation. The study was designed as a reproducible benchmark and as preparatory work toward future development of clinically meaningful H&E-based PARPi response models in locally curated HGSOC cohorts.

## Literature review

### HRD biology and therapeutic relevance in ovarian cancer

HGSOC is defined by extensive genomic instability, frequent TP53 mutation, and recurrent defects in homologous recombination repair.^1–4^ The discovery that BRCA1/2-deficient cancers are selectively sensitive to PARP inhibition provided the biological basis for synthetic lethality in ovarian cancer.^5,6^ Subsequent clinical trials established PARPi maintenance therapy as a major component of ovarian cancer treatment, particularly in BRCA-mutated and HRD-positive disease.^11–15^

The clinical assessment of HRD remains complex. BRCA1/2 testing identifies a highly relevant subset, but HRD can also arise through epigenetic silencing, including BRCA1 promoter methylation, or through alterations in other homologous recombination repair genes.^16–20^

Genomic scar approaches attempt to capture the cumulative consequences of defective DNA repair rather than single-gene status alone. LOH, TAI, and LST have been used individually or in composite HRD scores, but these assays are not uniformly available in public datasets and may not capture dynamic restoration of homologous recombination proficiency.^21–25^

### TCGA-OV as a public research resource

TCGA-OV remains one of the most important public molecular pathology datasets for HGSOC, providing genomic, transcriptomic, methylation, copy-number, and histopathological data.^1,26^ The dataset has enabled multiple downstream analyses of ovarian cancer biology, molecular subtypes, survival, and therapy-associated biomarkers. However, TCGA-OV was not designed as a modern PARPi-response dataset. Its public clinical and molecular data do not provide a harmonised set of validated clinical HRD assay calls, genomic scar scores, BRCA1 promoter methylation status across all relevant cases in a directly analysis-ready form, and PARPi response labels. This creates a label-quality problem for computational pathology studies using TCGA-OV.

### Computational pathology and molecular prediction from H&E

Deep learning has transformed computational pathology by enabling WSI-based prediction of diagnostic categories, molecular alterations, prognosis, and treatment response.^27–35^ Studies have shown that H&E morphology can be associated with EGFR mutation, microsatellite instability, tumour mutational burden, transcriptomic states, and survival outcomes across several cancers.^30–35^ In ovarian cancer, published studies have investigated BRCA mutation prediction, HRD prediction, survival, and molecular subtype inference from WSIs.^45–48^ These findings support the biological plausibility of morpho-molecular modelling but also highlight the need for careful validation, transparent labels, and patient-level separation.

### Multiple instance learning and leakage control

MIL is particularly suited to WSI analysis because each slide or patient can be represented as a bag of image tiles or tile-derived features, with only a slide-level or patient-level label available.^36–39^ Attention-based MIL provides a mechanism for aggregating patch-level information into slide-level or patient-level predictions and can support qualitative review through attention maps.^36,37^ However, WSI studies are vulnerable to data leakage when patches or slides from the same patient appear across training and test folds. Leakage can inflate performance and undermine reproducibility. For clinically meaningful computational pathology, splitting must occur at the patient level, especially when multiple slides are available from the same case.

### Foundation models in pathology

Large-scale histopathology foundation models have recently been introduced as general-purpose feature extractors for downstream computational pathology tasks.^40–44^ UNI is a vision foundation model trained on a large histopathology corpus, while CONCH is a vision-language pathology foundation model trained using image-caption pairs and histopathology text-image alignment.^40,41^ Virchow and other large-scale WSI models have further demonstrated the potential of pretraining on diverse histopathology data.^42–44^ Although these models can improve data efficiency and generalisation, they do not eliminate the need for high-quality labels, leakage-controlled evaluation, and disease-specific validation.

### Rationale for the present exploratory benchmark

The present study addresses a specific methodological gap: how should TCGA-OV be used responsibly for H&E-based HRD modelling when validated clinical HRD and PARPi-response labels are absent? Rather than presenting a clinical predictor, we frame TCGA-OV as a method-development and label-refinement benchmark. This allows systematic evaluation of encoder choice, label definition, leakage control, and performance limits under realistic public-data constraints. The work also provides an auditable preparatory framework for future modelling of PARPi response in clinically curated HGSOC cohorts.

## Materials and methods

### Study design

This was a computational pathology benchmarking study using public TCGA-OV H&E WSIs and publicly available molecular evidence. The objective was to evaluate whether H&E WSIs contain a measurable morphology-linked signal associated with HRD labels, and to determine whether label refinement and encoder choice affect model performance. The conceptual framework of this study is illustrated in Figure 1.

**Figure 1.**
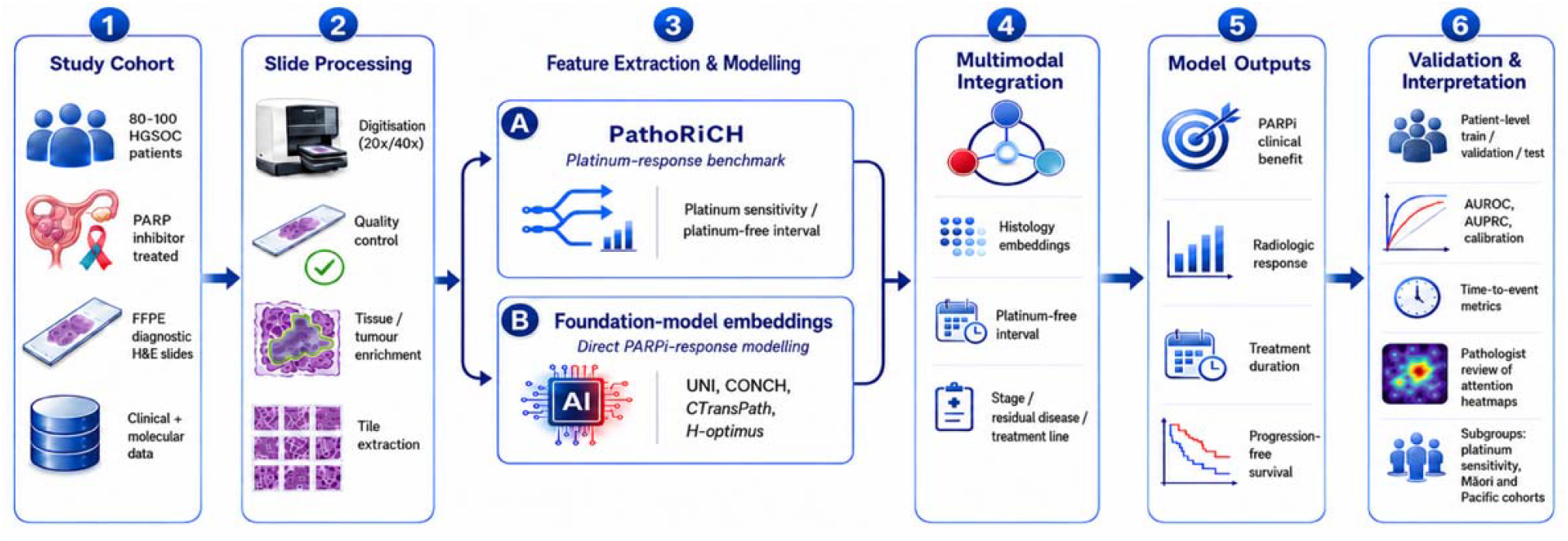
Conceptual framework overview.

### Cohort assembly

The initial public TCGA-OV context comprised 608 clinical cases. A total of 1,481 candidate pathology or slide files were identified. From these, 778 SVS files were downloaded and passed readability and quality-control checks. The main modelling cohort comprised 717 frozen-primary tissue-section/biospecimen slides from 316 patients. Diagnostic FFPE DX slides were excluded from model training and model selection because the 61 available DX slides had complete patient overlap with the frozen-primary cohort. DX slides were therefore reserved only as a potential future out-of-distribution slide-type analysis, requiring fold-aware routing.

### Label curation

Two HRD labels were constructed. The initial mutation-only molecular label classified patients as positive if they had BRCA1/2 or homologous recombination gene mutation evidence available through public molecular data. This produced 78 positive and 238 negative patients, corresponding to a positive prevalence of 24.7%.

The refined methylation-enhanced molecular label incorporated BRCA1 promoter methylation evidence in addition to mutation-based evidence. BRCA1 promoter methylation was derived from available methylation data for the 316 WSI patients using a primary beta-value threshold of >=0.30. This recovered 33 additional methylation-driven positive cases, resulting in 111 positive and 205 negative patients, corresponding to a positive prevalence of 35.1%.

Validated clinical HRD assay labels, genomic scar scores, LOH/TAI/LST composite scores, and PARPi/platinum response labels were not available in a form suitable for the present modelling workflow. These were therefore not inferred or imputed.

### Whole-slide image processing and feature extraction

All included slides underwent readability checking, tissue-mask assessment, patch-coordinate generation, and feature extraction. Six feature extractors were evaluated:

1. ResNet50, used as an ImageNet-pretrained baseline encoder, producing 2048-dimensional features.
2. UNI, used as a pathology foundation-model encoder, producing 1024-dimensional features.
3. CONCH, used as a pathology vision-language foundation-model encoder, producing 512-dimensional features.

Virchow2, used as a large-scale pathology foundation-model encoder, producing 2560-dimensional features.

Phikon-v2, used as a pathology foundation-model encoder, producing 1024-dimensional features.

UNI2-h, used as the strongest-performing pathology foundation-model encoder in this experiment, producing 1536-dimensional frozen tile features.

Encoder-specific HDF5 feature stores were generated for the frozen-primary slide set and used for patient-level modelling. DX slides were not included in feature-store training or model selection.

### Patient-level bag construction

The unit of inference was the patient. Features from available frozen-primary TS/BS slides were aggregated into patient-level bags. This design was selected to prevent slide-level leakage and to match the clinically relevant unit of inference. No random patch-level or slide-level train/test splitting was used.

### Model architecture

Patient-level attention-based multiple instance learning was used for binary classification. Each patient bag contained encoder-derived WSI features. The ABMIL aggregator produced patient-level probability estimates for each label definition. Separate models were trained for each encoder–label combination.

### Cross-validation and leakage control

All experiments used patient-level grouped 5-fold x 5-repeat stratified cross-validation, with 25 folds total. Grouping was performed by patient identifier. This ensured that no patient contributed slides or features to both training and test folds. Out-of-fold predictions were generated for all 316 patients. Bootstrap confidence intervals were calculated for AUROC and other key metrics where available. Validation-calibrated thresholds were used alongside default 0.5 thresholds for classification metrics.

### Evaluation metrics

Model performance was assessed using AUROC, AUPRC, no-skill baseline, balanced accuracy, sensitivity, specificity, F1 score, and Brier score where available. AUROC was treated as the primary discrimination metric, while AUPRC was interpreted in relation to the positive prevalence of each label definition. Because the initial mutation-only label and refined methylation-enhanced label had different class prevalences, raw AUPRC values were not directly compared without considering their no-skill baselines.

### Error analysis

High-confidence errors, borderline predictions, and calibrated-threshold classification outputs were reviewed for the best-performing HRD models. Methylation-driven positives recovered by the refined label were interpreted cautiously as improved molecular label curation rather than proof of a distinct methylation-specific morphology signal.

### Reproducibility and governance

The pipeline was designed with explicit acceptance gates: feature-store validation, label integrity checks, grouped cross-validation leakage checks, finite prediction verification, caveat compliance, and report generation. Reports documented input files, label versions, encoder type, feature dimensions, cross-validation design, patient counts, and model outputs. All model outputs were treated as research-use only.

## Results

### Cohort and slide inventory

The final modelling cohort consisted of 717 frozen-primary TCGA-OV WSIs from 316 patients. Encoder-specific feature extraction was performed for ResNet50, UNI, CONCH, Virchow2, Phikon-v2 and UNI2-h. DX slides were excluded from model selection because all DX patients overlapped with the frozen-primary cohort. This prevented the DX set from being misused as an independent validation set.

### Label refinement changed cohort composition

The initial mutation-only molecular label identified 78 positive and 238 negative patients. After incorporating BRCA1 promoter methylation, the refined methylation-enhanced molecular label identified 111 positive and 205 negative patients, with no unknown cases. Thus, label refinement increased the positive prevalence from 24.7% to 35.1% and recovered 33 additional methylation-driven positives.

### HRD-only encoder and label comparison

The HRD-only comparison showed modest but reproducible discrimination overall, with the strongest performance from UNI2-h under the refined methylation-enhanced molecular label.

Using the refined methylation-enhanced label, patient-level ABMIL with UNI2-h features achieved AUROC 0.634 (95% CI 0.571-0.698) and AUPRC 0.468 (95% CI 0.390-0.562), above the no-skill AUPRC/prevalence baseline of 0.351. At the calibrated threshold, the model achieved balanced accuracy 0.597 (95% CI 0.545-0.655), sensitivity 0.532 (95% CI 0.444-0.622), specificity 0.663 (95% CI 0.599-0.727), F1 score 0.494 (95% CI 0.417-0.568), and Brier score 0.233 (95% CI 0.222-0.244).

Comparative performance contextualised the UNI2-h result. UNI2-h with the initial mutation-only label achieved AUROC 0.628 and AUPRC 0.333. Refined-label Phikon-v2 and Virchow2 each achieved AUROC 0.582, with AUPRC 0.447 and 0.428, respectively. CONCH with the initial mutation-only label achieved AUROC 0.587, while ResNet50 with the refined label achieved AUROC 0.570. Clinical logistic-regression and random-forest baselines achieved AUROC 0.57 and 0.54, respectively.

Overall, the new HRD-only results shift the primary result from the earlier CONCH/ResNet50 framing to UNI2-h on the refined methylation-enhanced label. The result should be presented as proof-of-concept computational pathology evidence for a modest histology-associated molecular HRD signal, not as a clinical HRD assay, PARP-inhibitor response predictor, platinum-response predictor, or replacement for molecular testing.

**Table 1.** Best-model HRD prediction performance using UNI2-h with the refined methylation-enhanced molecular label.

| Metric | UNI2-h refined result | Bootstrap 95% CI / note |
| --- | --- | --- |
| AUROC | 0.634 | 0.571-0.698 |
| AUPRC | 0.468 | 0.390-0.562 |
| No-skill AUPRC / prevalence | 0.351 | Not applicable |
| Balanced accuracy | 0.597 | 0.545-0.655 |
| Sensitivity | 0.532 | 0.444-0.622 |
| Specificity | 0.663 | 0.599-0.727 |
| F1 score | 0.494 | 0.417-0.568 |
| Brier score | 0.233 | 0.222-0.244 |
| Confusion matrix | TN=136, FP=69, FN=52, TP=59 | Calibrated threshold |
| Calibrated threshold | 0.512 | Youden median |

**Table 2.** Comparative HRD-only model performance across selected encoders and clinical baselines.

| Model | Dim | Label | n | AUROC | AUPRC | Sens. | Spec. |
| --- | --- | --- | --- | --- | --- | --- | --- |
| ABMIL + UNI2-h | 1536 | Refined methylation-enhanced | 316 | 0.634 | 0.468 | 0.532 | 0.663 |
| ABMIL + UNI2-h | 1536 | Initial mutation-only | 316 | 0.628 | 0.333 | 0.538 | 0.588 |
| ABMIL + Phikon-v2 | 1024 | Refined methylation-enhanced | 316 | 0.582 | 0.447 | 0.530 | 0.610 |
| ABMIL + Virchow2 | 2560 | Refined methylation-enhanced | 316 | 0.582 | 0.428 | 0.560 | 0.580 |
| ABMIL + CONCH | 512 | Initial mutation-only | 316 | 0.587 | Not listed | 0.620 | 0.550 |
| ABMIL + ResNet50 | 2048 | Refined methylation-enhanced | 316 | 0.570 | 0.391 | 0.590 | 0.590 |
| Clinical baseline LR | Clinical | HRD baseline | 316 | 0.570 | Not listed | 0.590 | 0.480 |
| Clinical baseline RF | Clinical | HRD baseline | 316 | 0.540 | Not listed | 0.390 | 0.640 |

### Effect of refined methylation-enhanced label definition

The refined methylation-enhanced label altered cohort composition by increasing HRD positivity from 24.7% to 35.1%. This was methodologically important because it incorporated BRCA1 promoter methylation, a biologically relevant mechanism of BRCA pathway disruption, and changed the no-skill AUPRC baseline. The primary refined-label result should therefore be interpreted in relation to the higher positive prevalence rather than compared directly with mutation-only AUPRC values.

### Best-model classification profile

At the calibrated threshold of 0.512, the best UNI2-h refined-label model classified 136 patients as true negatives, 69 as false positives, 52 as false negatives, and 59 as true positives. This profile produced moderate specificity but only modest sensitivity, reinforcing that the model captures a research-grade signal rather than a clinically actionable classification rule.

### Label refinement interpretation

The 33 methylation-driven positives recovered by the refined label should be interpreted as improved molecular label curation rather than proof of a distinct methylation-specific histological phenotype. Attention or heatmap-based interpretation would require saved fold checkpoints, pathologist review, and external validation before biological claims are made.

### Heatmap and DX analysis status

Attention heatmaps were not generated for final interpretation because fold checkpoints were not saved. A set of priority qualitative cases should be reviewed after checkpoint-saving is implemented. DX out-of-distribution evaluation was also not performed because DX slides had complete patient overlap with the frozen-primary cohort and could not be used as an independent validation set.

## Discussion

This benchmarking study demonstrates that H&E WSIs from TCGA-OV contain a modest but reproducible morphology-linked signal associated with research-grade molecular HRD labels. The strongest result was obtained using patient-level ABMIL with UNI2-h features for the refined methylation-enhanced molecular label, achieving AUROC 0.634 and AUPRC 0.468. These results remain clearly below the level required for clinical deployment, but they are scientifically informative because they show how label definition, encoder selection, and leakage-controlled evaluation shape apparent performance in a public computational pathology dataset.

There are several learnings from this benchmarking. First, it presents a technically reproducible computational pathology pipeline. The workflow includes WSI ingestion, slide filtering, tissue-mask and patch-coordinate processing, encoder-specific feature extraction, patient-level bag construction, ABMIL modelling, grouped cross-validation, bootstrap confidence intervals, and structured reporting. This end-to-end design bears significance because computational pathology studies often fail not because of model architecture alone, but because of incomplete provenance, inconsistent labels, uncontrolled slide inclusion, or leakage between training and test data. Second, the study provides a comparative benchmark across a conventional ImageNet baseline, multiple pathology foundation-model encoders, and clinical baselines. Foundation models are increasingly used as feature extractors in computational pathology, but their performance is task-dependent and label-dependent. The updated results show that UNI2-h performed best under the refined methylation-enhanced molecular label, while Phikon-v2, Virchow2, CONCH, ResNet50, and clinical baselines showed lower discrimination. This supports the broader lesson that foundation-model pretraining can help but does not overcome weak, incomplete, or research-grade labels. Third, the study contributes a label-refinement analysis. The refined methylation-enhanced molecular label added BRCA1 promoter methylation to a mutation-only HRD label and recovered 33 additional positives. This is biologically relevant because BRCA1 promoter methylation is an established mechanism of BRCA pathway disruption in ovarian cancer. The study shows that label refinement can alter class prevalence, AUPRC baseline, model discrimination, and error profile. This makes the label-curation component a central methodological contribution rather than a minor preprocessing step. Fourth, the study offers an honest modest-signal and negative-learning story. In a field where computational pathology results may be overstated, this study provides a transparent example of what can and cannot be inferred from TCGA-OV. The best AUROC of 0.634 is informative but not clinical-grade. It indicates that morphology may encode some HRD-associated molecular information, but also that public labels and cohort constraints limit clinical interpretability. Such rigorous modest-signal benchmarking is valuable because it helps prevent premature claims and guides future study design. Fifth, the study creates a translational bridge to a clinically meaningful next phase. The authors performed this work as exploratory preparation for developing a predictive model of PARPi response from H&E images. The current TCGA-OV benchmark establishes the pipeline, label-governance principles, encoder framework, leakage-control strategy, and reporting infrastructure. The next scientific step is not to over-interpret TCGA, but to apply the same framework to an HGSOC cohort with curated PARPi responder/non-responder labels, treatment duration, progression-free survival, platinum response, molecular HRD data, and pathologist-reviewed qualitative outputs.

Reproducibility of the pipeline is a major strength of the present study. The pipeline used explicitly versioned feature stores, label namespaces, patient identifiers, slide-type filters, and grouped cross-validation. Each encoder-specific feature store was validated for slide count, feature dimensionality, finite values, and gate-readiness. The study also maintained separation between the initial mutation-only label and the refined methylation-enhanced label, preventing metrics from being blended across label definitions. The patient-as-bag design is particularly important. In WSI datasets, a single patient may contribute multiple slides, tissue sections, or patch sets. If random patch-level or slide-level splitting is used, a model may effectively see tissue from the same patient during both training and testing, leading to inflated performance. By grouping all folds by patient identifier, the present study reduced this risk and made the reported AUROCs more credible. The study also used repeated grouped stratified cross-validation rather than a single arbitrary split. This is appropriate for a modest-sized cohort of 316 patients, where model performance may be sensitive to fold composition. Bootstrap confidence intervals further support transparent interpretation of uncertainty. The fact that several confidence intervals crossed chance performance was not hidden; rather, it was incorporated into the biological and methodological interpretation.

An important limitation of this study is label quality. The model does not predict a validated clinical HRD assay label. It does not predict a genomic scar score. It predicts only a research-grade molecular HRD label derived from public molecular evidence. This distinction is essential. HRD is not equivalent to BRCA mutation status alone. It can reflect BRCA1/2 mutation, BRCA1 promoter methylation, alterations in other homologous recombination repair genes, genomic scar patterns, and potentially functional DNA repair states. Conversely, a tumour may carry historical evidence of HRD but regain homologous recombination proficiency through reversion mutations or other resistance mechanisms. Therefore, mutation-only and methylation-enhanced labels are biologically incomplete.

The refined methylation-enhanced label improved biological plausibility by incorporating BRCA1 promoter methylation, but it still lacked validated LOH/TAI/LST genomic scar scoring. It may therefore miss scar-positive tumours without identifiable mutation or methylation evidence. It may also include cases where molecular evidence does not translate into current functional HRD. These limitations likely contribute to modest AUROC values, imperfect sensitivity and specificity, and the need for external validation.

TCGA-OV also lacks a clinically useful PARPi-response endpoint for this modelling task. Many TCGA cases predate contemporary PARPi maintenance therapy, and clinical treatment annotations are not sufficiently harmonised for direct response modelling. Therefore, TCGA-OV is best treated as a public method-development and label-refinement benchmark rather than a definitive response-prediction dataset.

The observed AUROC of 0.634 for the best UNI2-h refined-label model suggests a modest histology-linked signal rather than robust clinical prediction. The result is stronger than the earlier CONCH and ResNet50 framing, but the discrimination remains insufficient for clinical use. The updated comparison also indicates that encoder choice matters: UNI2-h was strongest in this internal experiment, while Phikon-v2, Virchow2, CONCH, ResNet50, and clinical baselines were lower-performing comparators.

AUPRC interpretation also requires caution. Because the refined methylation-enhanced label increased positive prevalence from 24.7% to 35.1%, raw AUPRC values cannot be compared directly without accounting for the no-skill baseline. The best refined-label AUPRC of 0.468 should be interpreted against the 0.351 prevalence baseline and not as evidence of clinical-grade prediction.

This study should be viewed as preparation for the clinically important question: can H&E morphology help predict PARPi response in HGSOC? The present results do not answer that question directly, but they provide the infrastructure required to pursue it rigorously. A future PARPi-response study by authors will include a clinically curated cohort of patients treated with PARPi, with response/non-response labels defined using progression-free survival, duration of clinical benefit, radiological response, treatment line, platinum sensitivity, HRD assay status, BRCA status, residual disease, stage, and other relevant covariates. This future study will compare morphology-only prediction against clinical-only and molecular-only baselines, then test whether fusion models add incremental value. It will also undertake external validation, calibration, decision-curve analysis, and pathologist review of attention-selected regions.

### Limitations

This study has several limitations. First, the labels are research-grade molecular proxies rather than validated clinical HRD assay labels. Second, no public genomic scar score was available in the implemented workflow, preventing direct modelling of LOH/TAI/LST-defined HRD. Third, TCGA-OV does not provide a suitable PARPi-response or platinum-response target for contemporary clinical prediction. Fourth, the cohort size was modest, with 316 patients, and the best AUROC of 0.634 indicates weak-to-moderate rather than clinical-grade discrimination. Fifth, DX slides could not be used as an independent validation set because of complete patient overlap with the frozen-primary cohort. Sixth, heatmaps were deferred because fold checkpoints were not saved and would require pathologist review. Finally, the study used feature-extraction-based modelling rather than end-to-end WSI training; while appropriate for reproducible benchmarking, this may limit representation learning for subtle phenotype discovery.

## Conclusion

This exploratory computational pathology study shows that TCGA-OV H&E WSIs contain a modest but reproducible signal associated with research-grade molecular HRD labels. Label refinement using BRCA1 promoter methylation altered cohort composition, increased positive prevalence, and enabled the strongest internal result with UNI2-h features, but did not produce a clinically deployable model. Foundation-model benchmarking showed task-dependent performance, with UNI2-h performing best under the refined label and other encoders or clinical baselines performing less strongly. The study is publishable as a reproducible, leakage-controlled methods and benchmarking paper, provided that claims remain focused on research-grade molecular proxy prediction rather than clinical HRD diagnosis or PARPi/platinum-response prediction. Its principal value lies in establishing a transparent pipeline and label-governance framework for future H&E-based PARPi-response modelling in clinically curated HGSOC cohorts.

## Declarations

### Ethics approval and consent to participate

This study used publicly available, de-identified TCGA-OV data.

### Consent for publication

Not applicable.

### Availability of data and materials

TCGA-OV data are available through public repositories including the Genomic Data Commons and associated TCGA/cBioPortal resources. Derived manifests, label files, feature extraction scripts, and model reports can be made available.

### Authors’ contributions

Conceptualization: EU, MS, NS, Writing – Original Draft: EU, MS, Writing – Review & Editing: MS, NS; Supervision: EU

### Data Availability

Data is available from corresponding author on reasonable request.

